# ASMS: finding allele specific methylation in human genomes without phasing

**DOI:** 10.1101/2024.12.18.629129

**Authors:** Emanuele Raineri, Miguel Ángel Esteve Marco, Anna Esteve Codina

## Abstract

**Motivation:** Allele-specific methylation (ASM) refers to differential DNA methylation patterns between two alleles at a given locus. This phenomenon is often driven by genetic variants, such as single nucleotide variants (SNVs), which influence methylation in *cis* by affecting transcription factor or methylation regulator binding, leading to allele-specific differences. Understanding ASM is critical for elucidating gene regulation, as it impacts gene expression and contributes to normal biological variation as well as disease processes, including cancer [1] and autoimmune disorders [2],[3].

Another key driver of ASM is genomic imprinting, an epigenetic mechanism in which gene expression is regulated in a parent-of-origin-specific manner. Imprinted regions, marked during gametogenesis and maintained through cell divisions, are essential for growth, development, and metabolism. Dysregulation of imprinting is associated with developmental and metabolic disorders, such as Prader-Willi and Angelman syndromes, and certain cancers.

Detecting ASM across the genome remains challenging due to its tissue- and cell-specific nature and the technical difficulty of phasing reads to assign methylation patterns to specific alleles. Current ASM detection pipelines (e.g., [4])) often require phasing via genetic variants, a computationally intensive process that is limited in regions with low heterozygosity.

**Results:** To address these limitations, we developed asms (Allele-Specific Methylation Scanner), a tool designed to detect ASM directly from methylation data without the need for prior phasing. asms offers a faster and complementary approach to uncovering the regulatory effects of ASM, particularly in genomic regions where genetic variants or imprinting play a critical role.

For demonstration purposes we leverage the fact that reads generated by Oxford Nanopore (ONT) technology measure sequence and methylation status at once, but the same software can be used with other sequencing technologies.

`asms` can check thousands of loci in a short time. The initial list of loci to examine can be given by the user, or generated by `asms` through a genomic scan. The `asms cluster` subcommand separates the reads based on methylation.

If phasing results are available, `asms` can use them to verify whether distinct alleles correspond to distinct base modifications patterns. We benchmark our software using publicly available Ashkenazi trio data [5].

**Implementation and availability:** `asms` is implemented in `rust` and `python`. The software is available at https://github.com/ecmra/asms.

## Introduction

Allele specific methylation (ASM) can vary across tissues and individuals and can offer insights for the interpretation of GWAS results([6]) and of regulatory events in cancer ([7]). This paper presents a pipeline to detect allele specific methylation patterns which does not need a previous phasing step. This is the main difference between `asms` and already existing software that is designed to detect ASM, for example hap-ASM[4], Methphaser[8] (which uses methylation data to improve phasing), and Nanopore’s `modkit`.

The approach we take extends the analysis done in [9] (which runs a likelihood ratio test on short reads) to long reads and builds upon our work in [10] where we have shown that an EM algorithm can group long reads depending on their methylation patterns. Here we present an optimized implementation so that it is possible to conduct a genome wide study in a under an hour.

There are no other methods that we are aware of to study the heterogeneity of a set of genomic regions without phasing first; `asms` can also scan the genome and look at variant information to produce a first list of interesting loci. Reads with different methylation levels mapping to the same locus can also result from the presence of a mixture of cells in the sequencing sample: our software can tackle analyses where changes in methylation cannot be captured by phasing, e.g. as those happening across genetically identical tissues or cell types; in such cases methods based on haplotypes are irrelevant, but read clustering still applies.

## Methods

A typical workflow with `asms` is depicted in figure S1. The process begins by identifying intermediately methylated regions, which are regions likely to exhibit allele-specific methylation. Users can then cluster reads at these loci, with an option to filter the resulting list of clusters and perform a permutation test.

`asms` works on BAM files, assuming that they contain methylation information encoded in the `MM` tag as explained in the SAM specifications [11]; it has 4 main subcommands: `scan`, `scan-vcf`, `cluster`, and `filter`.

`scan` finds intermediately methylated (IM) blocks and it has options to specify the size of the blocks which are selected; scanning a human chromosome takes around 1 second; it builds a list of intermediate methylation blocks by selecting runs of at least 3 consecutive CpGs with methylation between 0.4 and 0.6 (by default: users can choose different lower and upper bounds). IM blocks tend to appear in clumps, hence we merge nearby (default 1000 bp, configurable) blocks into larger ones.

`scan-vcf` checks a list of SNPs/SNVs and selects those loci around heterozygous variants which have intermediate methylation. By default it looks at methylation in a window of plus or minus 500 bp from the variant. In our tests we use a list of SNPs generated with `Clair3`[12].

The `cluster` subcommand tries to group reads mapping to a locus into 2 clusters representative of different methylation states (fig S4). This is done via an EM algorithm based on a multivariate Bernoulli model using a faster variant of the `cvlr` software [10]. This subcommand runs on a list of loci and outputs a file containing summary statistics for each region; while the first scanning step is used to build a list of potentially interesting regions, the `cluster` subcommand helps distinguishing unstructured intermediate methylation from regular patterns appearing on different reads. The initial point of the EM optimization is chosen non deterministically; this is not uncommon in clustering and to make the paper reproducible we use the option `--seed 7`.

The subcommand `filter` selects a subset of clusters by looking at their summary statistics and according to some simple criteria in order to define putative allele-specific methylation (ASM) regions; since all the intermediates results are stored in BED format, it is easy to select regions according to other criteria; at the moment `filter` uses the following (configurable) thresholds:

- total number of reads being clustered must be greater or equal to 10,
- they must cover a minimum of 2 CpGs,
- each cluster cannot contain less than 20% or more than 80% of all the reads and
- the median absolute difference in methylation must be greater or equal to 0.2.

## Results

In this section we evaluate the `asms` workflow and where possible we compare with allele specific methylation obtained by SNV calling and phasing. We use the publicly availabe Ashkenazi samples described in [5]. Specifically we look at `chr15` of sample `HG002`; this corresponds to a BAM file containing 529743 reads with quality above 10, of which 264118 are phased; reads are also augmented with methylation measurements stored in `MM` tags.

For comparison, we use the publicly available phasing data to look for allele specific methylation by running `modkit dmr pair --segment` (with default settings) to find out regions which are methylated differently across haplotypes. The command line we used for `modkit` is in the supplementary materials. `modkit` finds 25663 segments which are differentially methylated across haplotypes; those are typically much shorter than the 2055 merged IM blocks found by `asms` (fig S2); infact roughly half of the segments selected by `modkit` contain 1 CpG.

We then cluster the regions we found after scanning, and filter them through `asms filter`. We obtain 1105 ASM regions, 248 of which have no overlap with the output of `modkit dmr`. Since both `asms` and `modkit` effectively divide the reads over a locus in two groups (clusters), we look at the difference in methylation across the clusters in fig 1(A). Groups created by `asms` tend to differ more in methylation probably because `modkit` does not filter at all on methylation values when partioning the reads. We then look specifically at the regions found by `asms` and not by `modkit`; they have intermediate methylation levels (fig S3) and are mostly covered by haplotagged reads; there are 24 of them though, that are covered only by non-haplotagged reads; in those cases `asms` can still build methylation clusters whereas of course the pipeline based on `modkit` cannot draw any conclusion.

**Figure 1:**
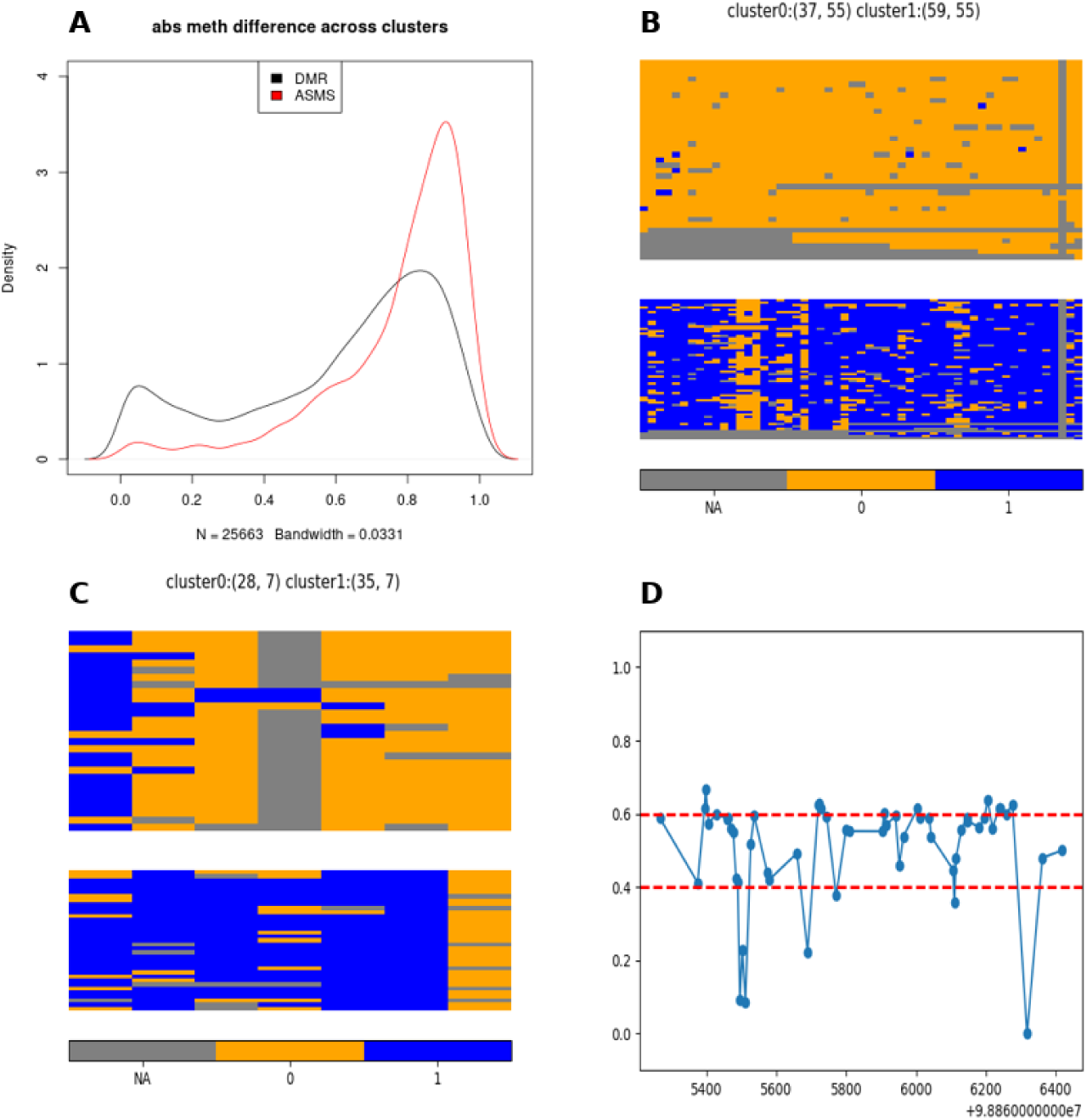
(A)difference in methylation across clusters defined by asms and modkit. asms tends to highlight regions with starker differences across read clusters. (B)clustering of the reads in the IGF1R region. (C)reads cluster around chr15:101492101 C>T (genotype is 1|0). (D)methylation in ICR region IGF1R(chr15:98865267-98866421).

In order to explore the significance of ASM regions we checked whether they overlap with promoters and we obtain that 144 intersect a promoter region on chr15 (defined as from 1000 bases upstream to 500 bases downstream of the transcription start site) at least once. By generating 10000 sets of random regions with the same length as the true ones we get a maximum of 102 intersection with promoters, hence the empirical p-value of our overlap is less than 10*^−^*^4^. Further enrichment with other interesting genomic landmarks (ehnancers, introns, etc…) can be easily computed in a similar way.

We also ran the clustering algorithm on the annotated [13] ICR regions on chr15. In this case since we are considering only 20 loci we can run a permutation test (with `asms pval`) to assign a p-value to the clusters; `asms` finds that 16 regions have p-value less than 10*^−^*^3^. Some ICR regions: SNORD115Cluster (`chr15:25181244-25184030`), and SNORD116Cluster (`chr15:25051256-25056714`, `chr15:25073965-25093073`) seem to be highly variable and not statistically significant in our approach; using phasing we can not find a partition either. We conclude these are regions with disordered methylation which are inherently difficult to partition automatically (See figures in supplementary materials). It’s easy to distinguish regions where the clustering did not work because one cluster has a small size compared to the other which contains almost all reads; in the case of imprinting control regions we expect clusters to have the same size. There are 3 regions, among the ones which we find to be significant which also have skewed cluster populations. These are TUBGCP5 (`chr15:23039853-23039919`), SORD (`chr15:45022590-45022746`) and PWARSN,SNORD107,PWARSN,PWAR5 (`chr15:24981888-24985149`); the methylation there is not intermediate. We also compared the clustering computed by `asms` with the haplotype tags generated by `Clair3` on the same regions and we observe that the partitions obtained by haplotagging is almost always significantly correlated with the partitions computed by `asms` (see table 1 in Supplementary material).

Finally, we look at heterozygous variants selecting those which corresponds to stretches with intermediate methylation. After clustering and filtering we find 642 putative ASM regions around heterozygous variants; 283 of them do not intersect with any of the DMRs found by modkit (see also fig S25.) Here below in fig. 1(C) we see an example of one such ASM region.

## Discussion

Allele-specific methylation is an important regulatory marker, which can be caused by (among other things) genetic variants in *cis* and parent of origin effects. The most direct way of finding ASM regions consists in calling variants from genomic data, phasing the aligned reads and computing differential methylation across haplotypes. This strategy has some limitations. First of all calling variants and phasing is a slow process; second many loci are difficult to phase because of lack of heterozygous variants or sequencing artifact.

Our software uses a complementary approach: clustering reads only based on methylation measurements. It builds upon the concepts tested in [10] but adds substantial improvements. First it’s faster: its speed allows users to explore thousands of genomic regions. Second it adds the possibility to scan the genome either with no prior information or in correspondence to pre specified loci; third, `asms` automatically produces statistical summary of all the genomic stretches considered in the analysis and optionally can filter the results and compute p-values based on permutation testing.

Overall, integrating `asms` with traditional phasing-based pipelines can provide a more comprehensive understanding of methylation. Furthermore, once one has a list of variants it is possible to use `asms` to check for heterogeneous methylation at those loci very quickly. As highlighted above, `asms` also work in the case where heterogeneity is due to mixture of different cells.

We think this an useful tool to improve analyses based on haplotypes or when a sample contains cells with different methylation states.

## Supplementary materials

### pipeline

**Figure S1:**
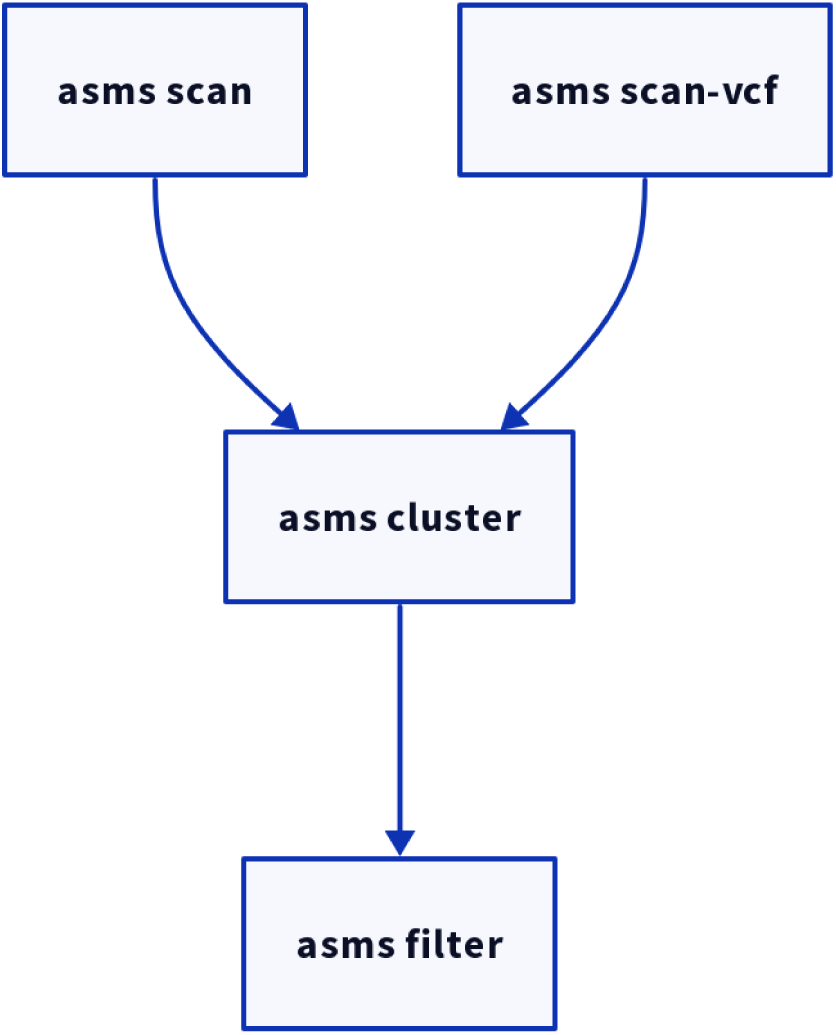
Typical workflow with `asms`. The first step is selecting potential ASM regions either by looking at methylation data or using genetic variants. These regions can be clustered and filtered. `asms` has a subcommand (`asms pval`) to assess the significance of the clustering via a permutation test.

### modkit script

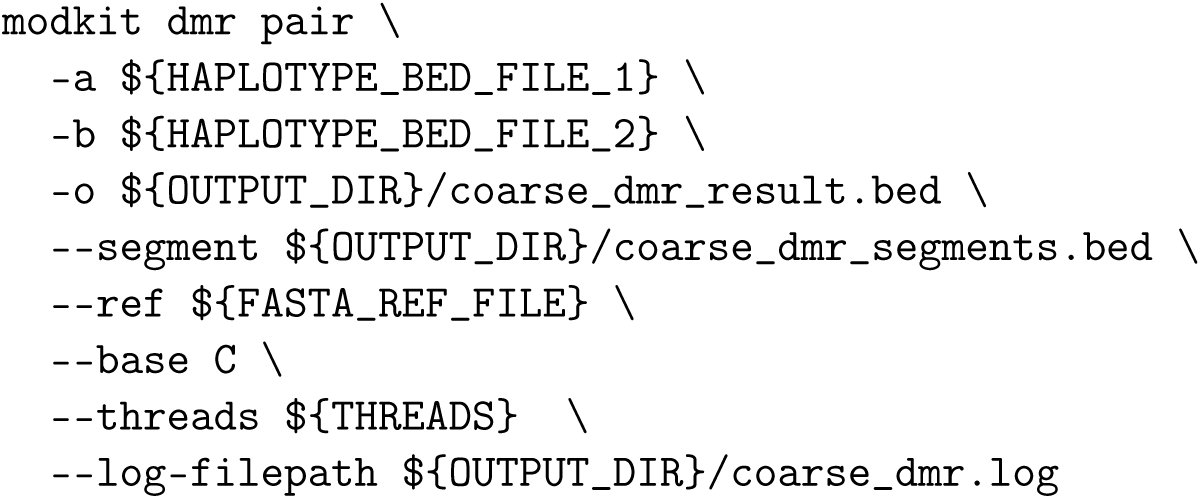

### modkit DMR and intermediate methylation blocks

**Figure S2:**
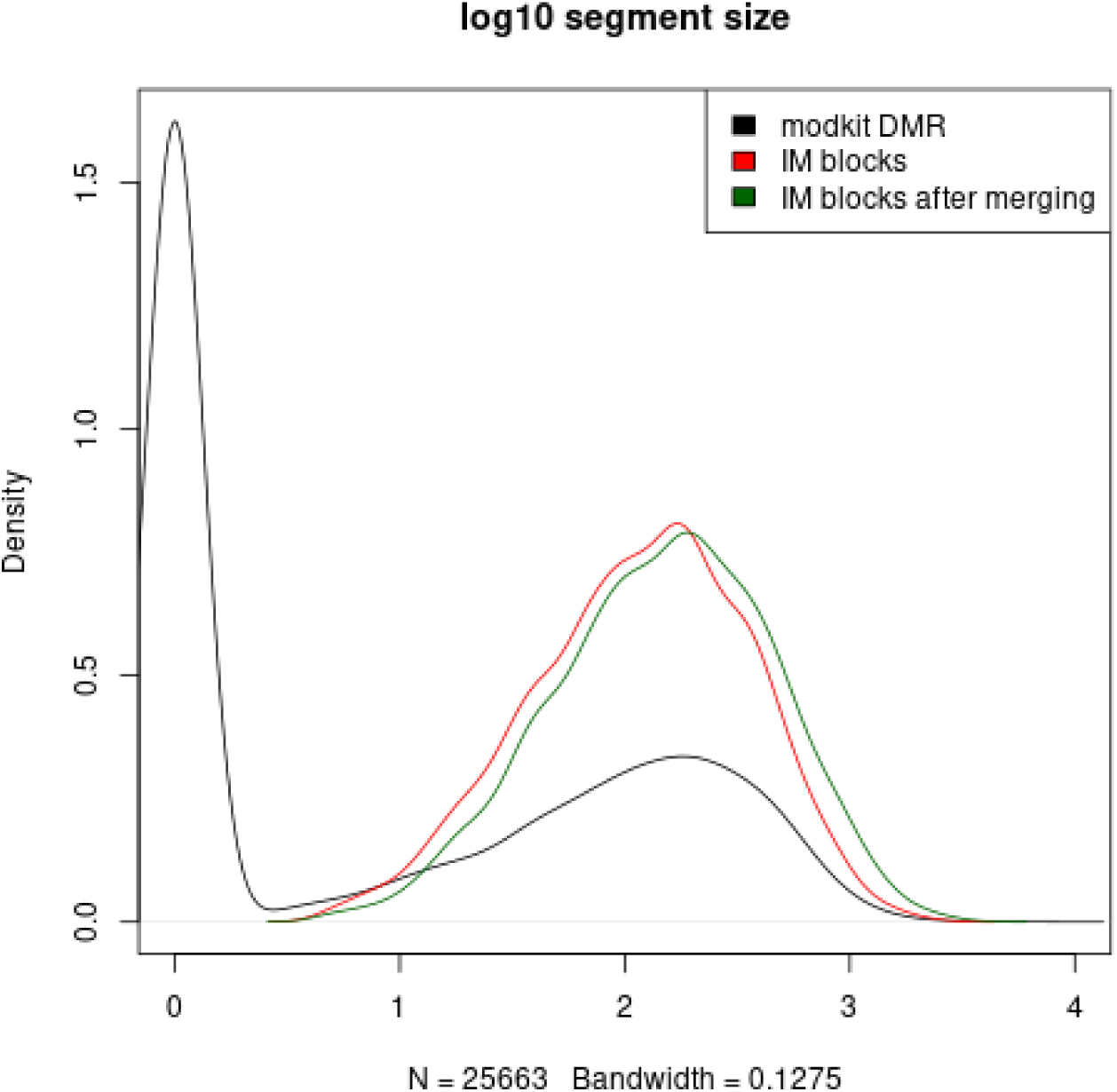
length distribution of modkit DMR segments and asms intermediate methylation regions. red not merged, dark green merged

**Figure S3:**
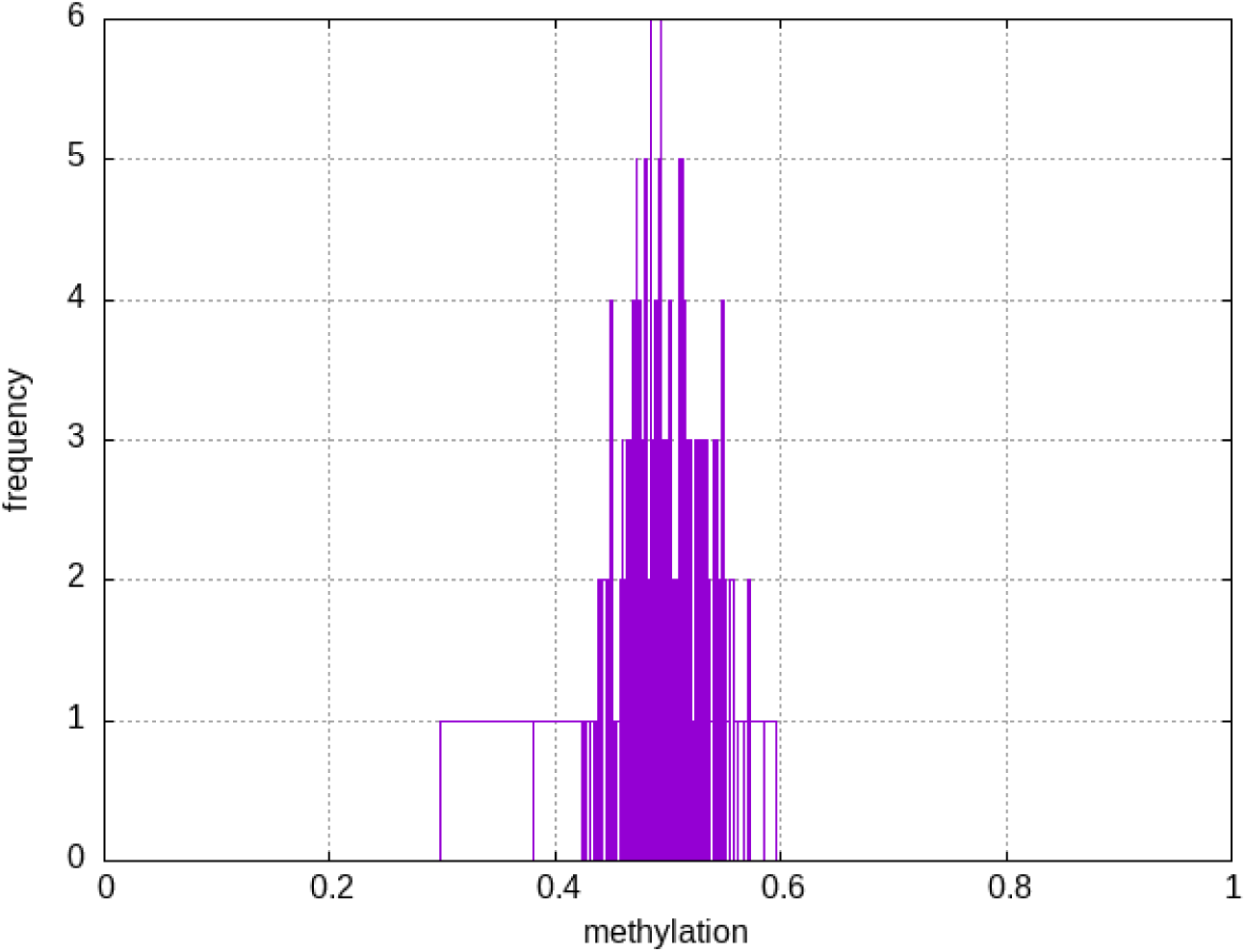
Average methylation at loci selected by `asms` and not found by modkit dmr These loci are typically covered by haplotagged reads, see main text.

### clustering algorithm

**Figure S4:**
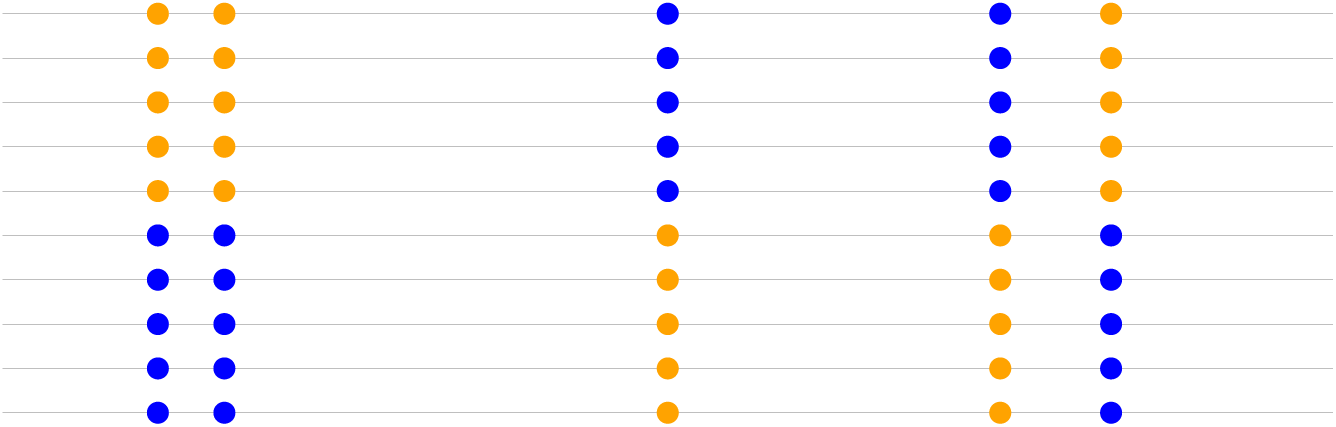
a cartoon of reads clustered according to methylation patterns. here blue=methylated, orange=unmethylated

### ICR regions

**Figure S5:**
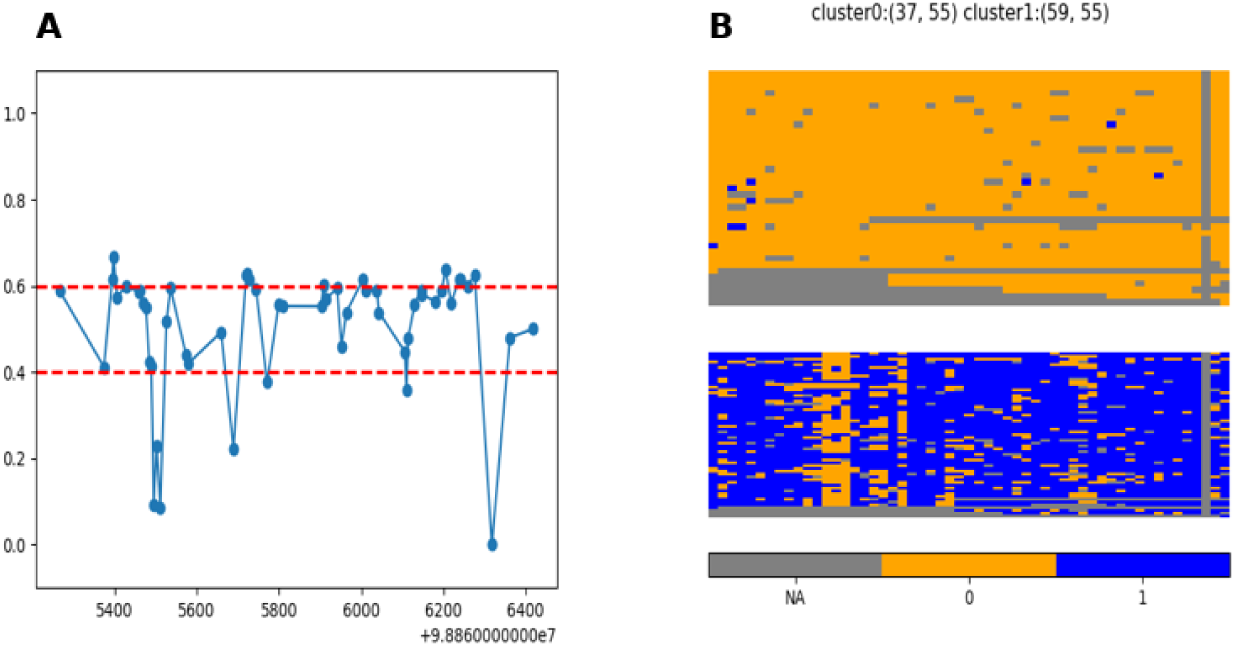
methylation in ICR region IGF1R(chr15:98865267-98866421) (A)methylation levels (B) clustered reads

**Figure S6:**
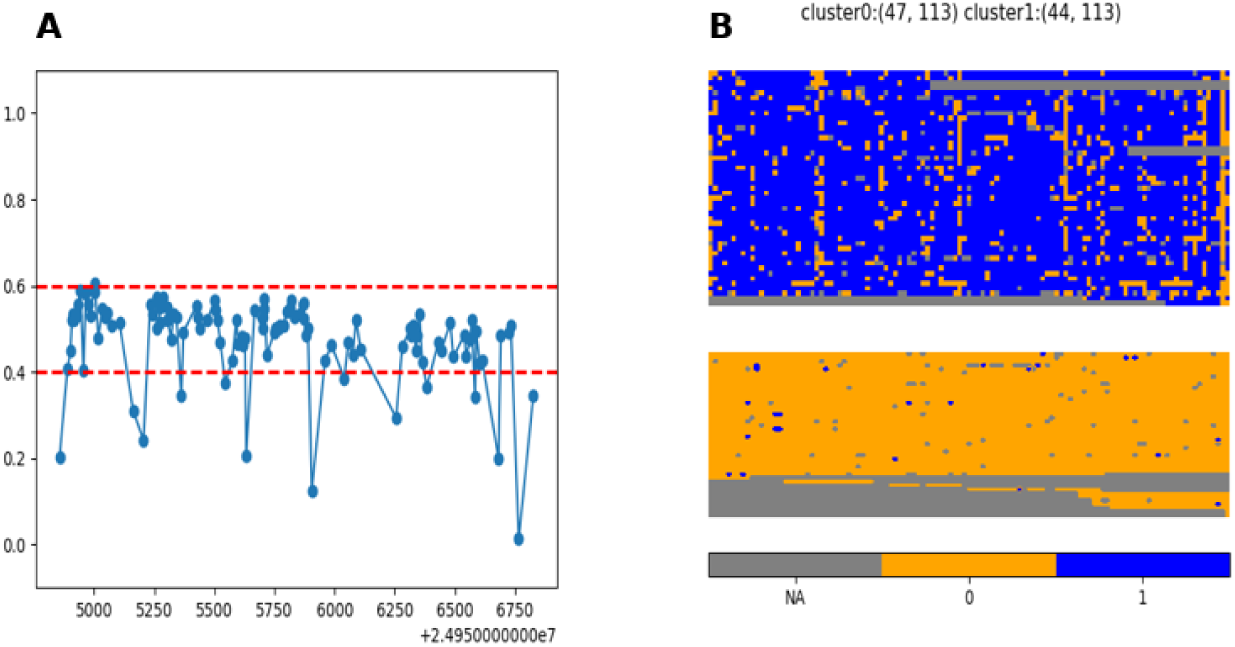
methylation in ICR region SNURF(chr15:24954857-24956829) (A)methylation levels (B) clustered reads

**Figure S7:**
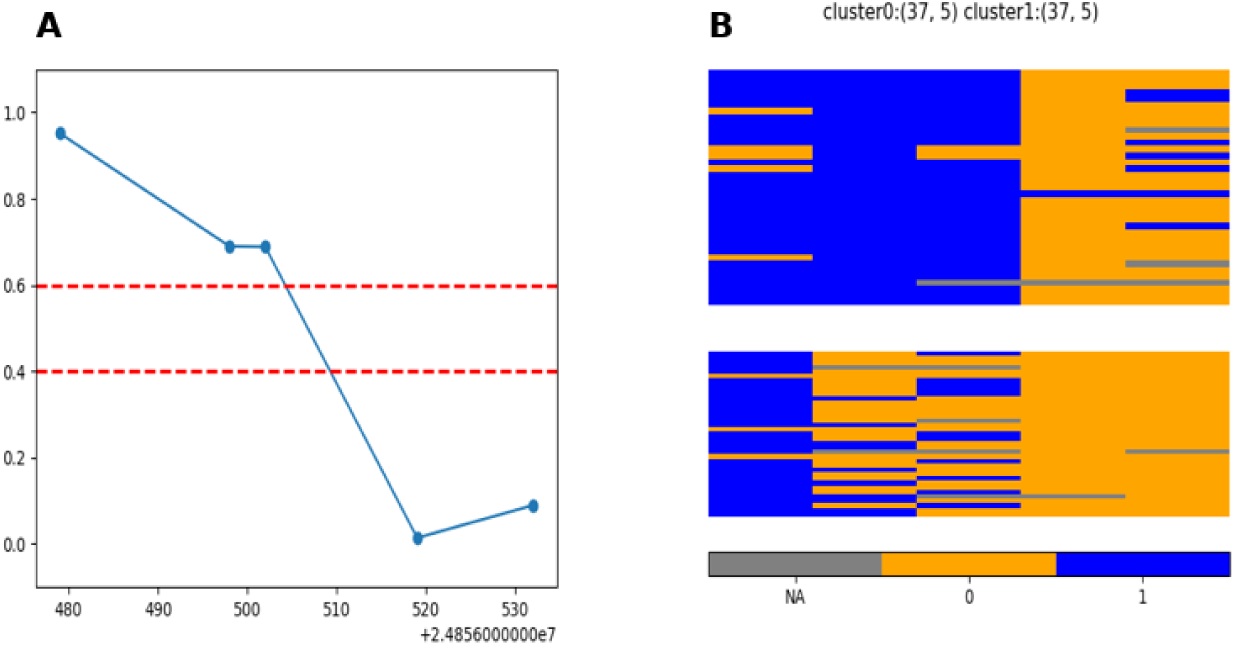
methylation in ICR region SNRPN(chr15:24856479-24856534) (A)methylation levels (B) clustered reads

**Figure S8:**
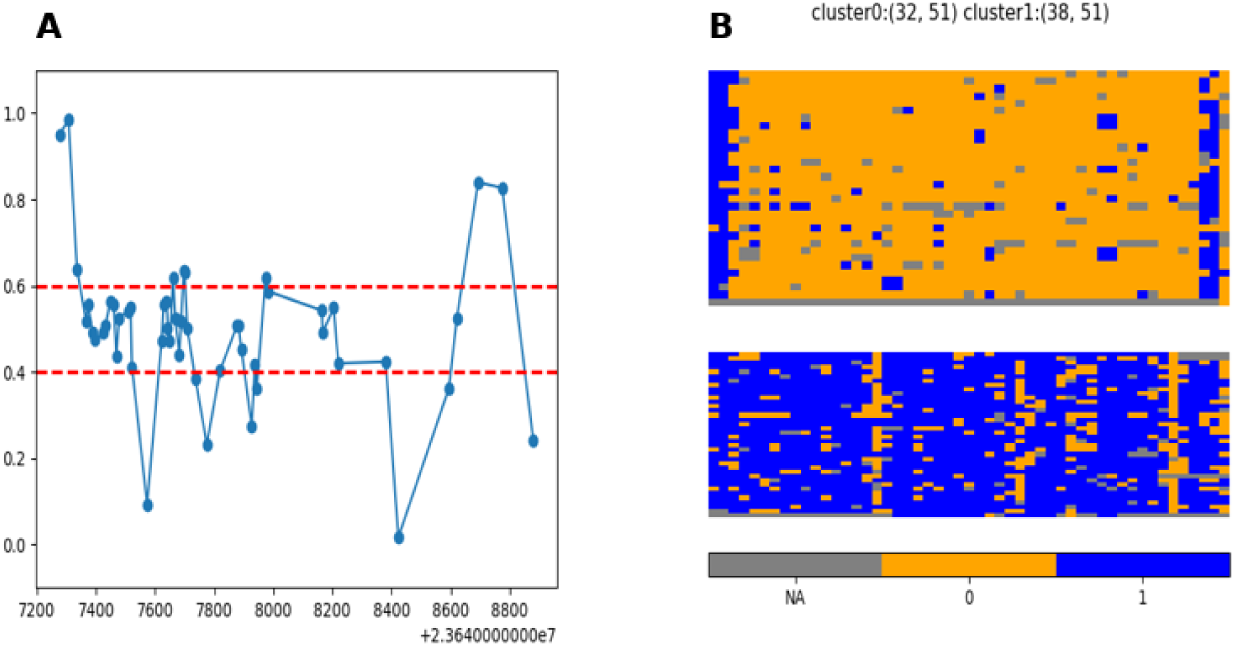
methylation in ICR region MAGEL2(chr15:23647278-23648882) (A)methylation levels (B) clustered reads

**Figure S9:**
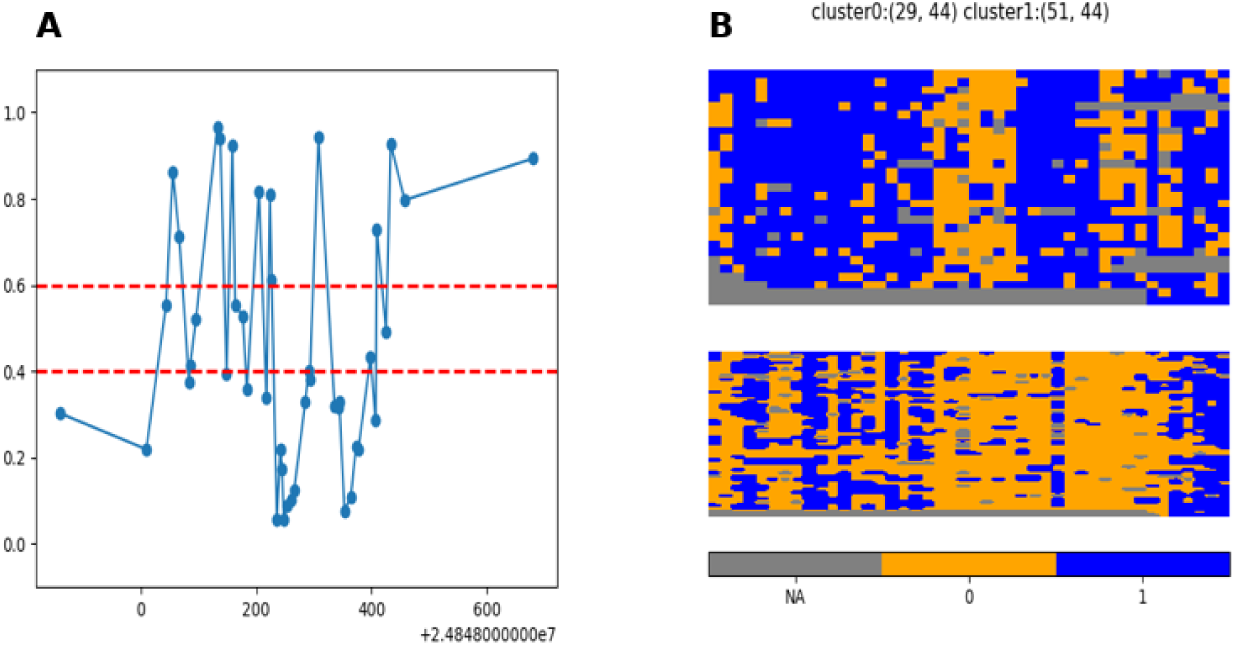
methylation in ICR region SNRPN(chr15:24847861-24848682) (A)methylation levels (B) clustered reads

**Figure S10:**
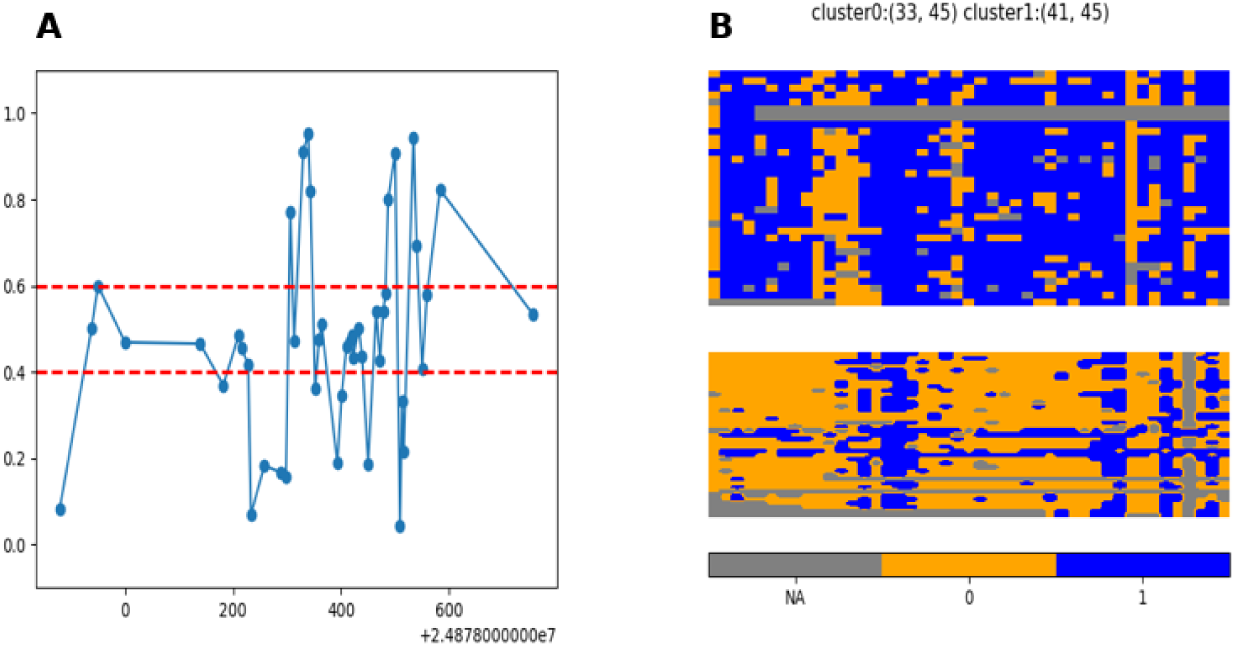
methylation in ICR region SNRPN(chr15:24877880-24878758) (A)methylation levels (B) clustered reads

**Figure S11:**
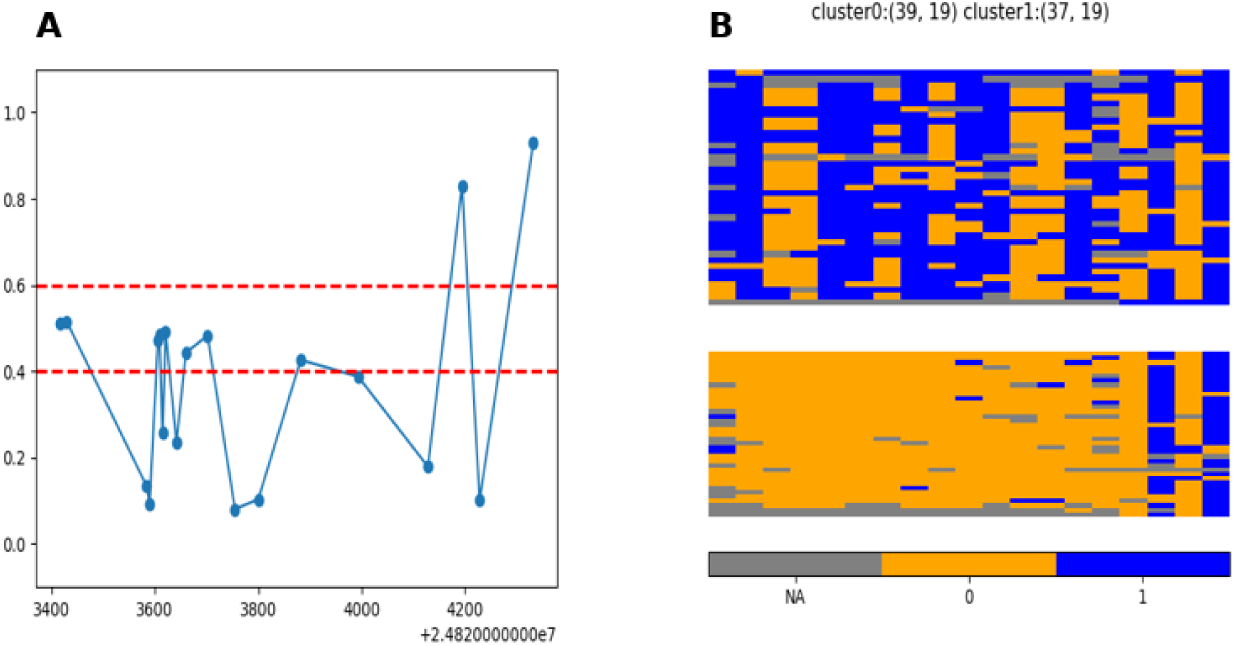
methylation in ICR region SNRPN(chr15:24823417-24824334) (A)methylation levels (B) clustered reads

**Figure S12:**
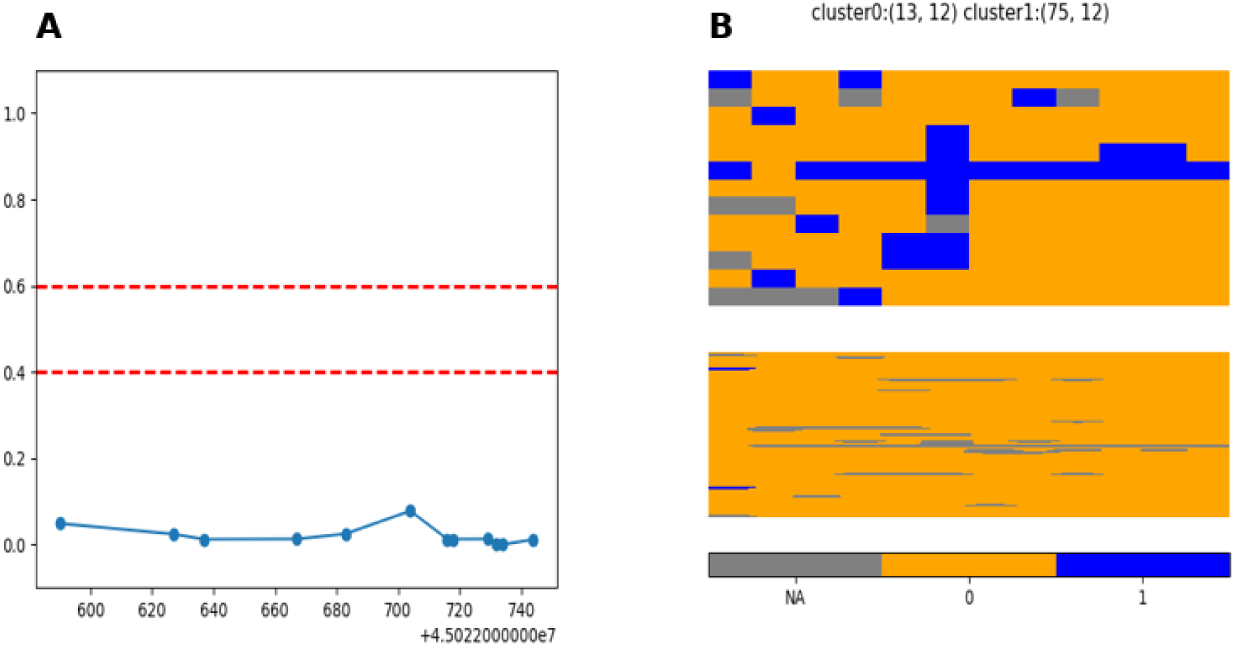
methylation in ICR region SORD(chr15:45022590-45022746) (A)methylation levels (B) clustered reads

**Figure S13:**
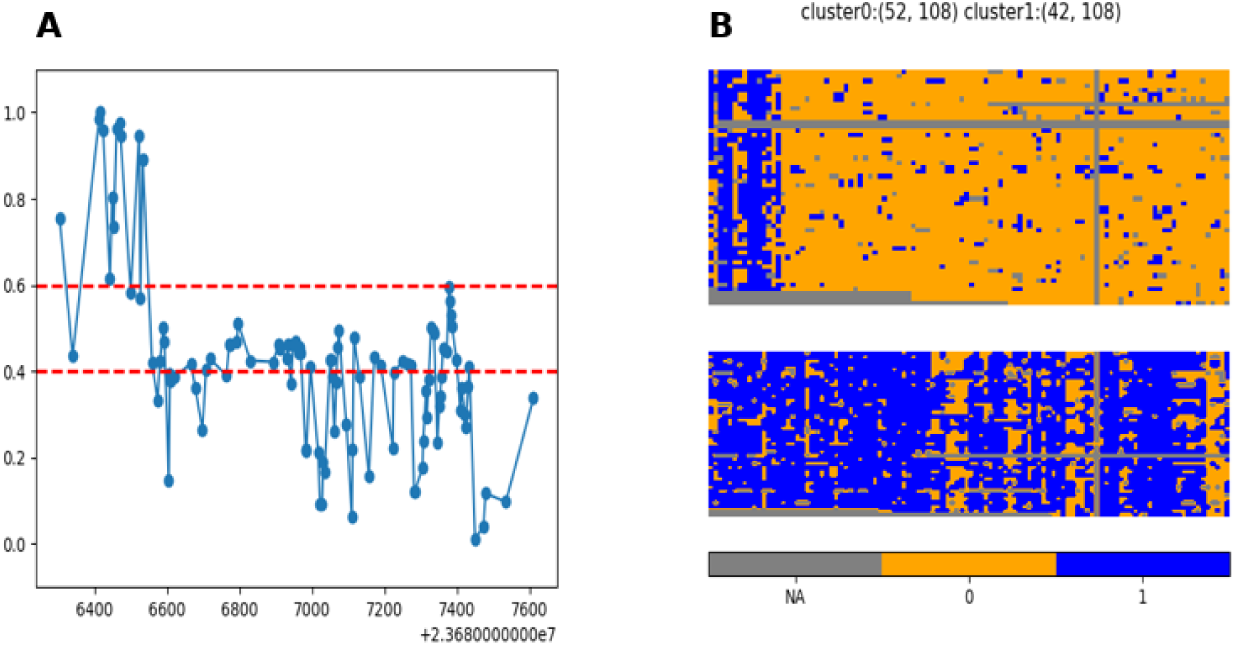
methylation in ICR region NDN(chr15:23686304-23687612) (A)methylation levels (B) clustered reads

**Figure S14:**
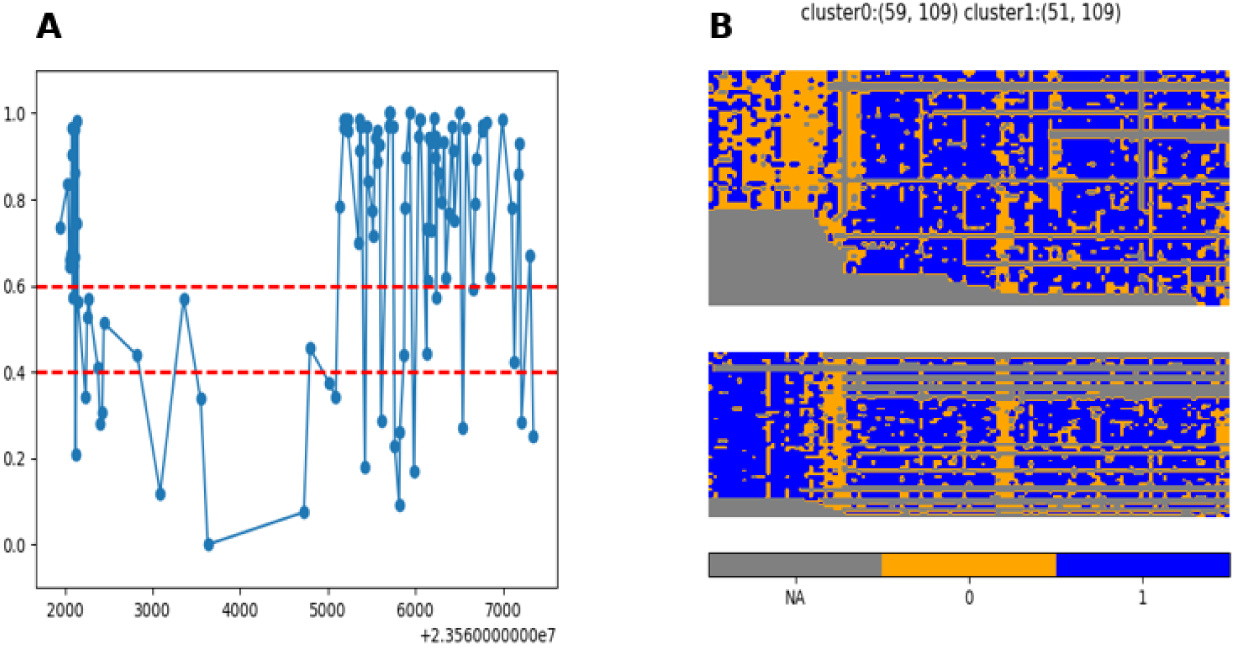
methylation in ICR region MKRN3,MIR4508(chr15:23561939-23567348) (A)methylation levels (B) clustered reads

**Figure S15:**
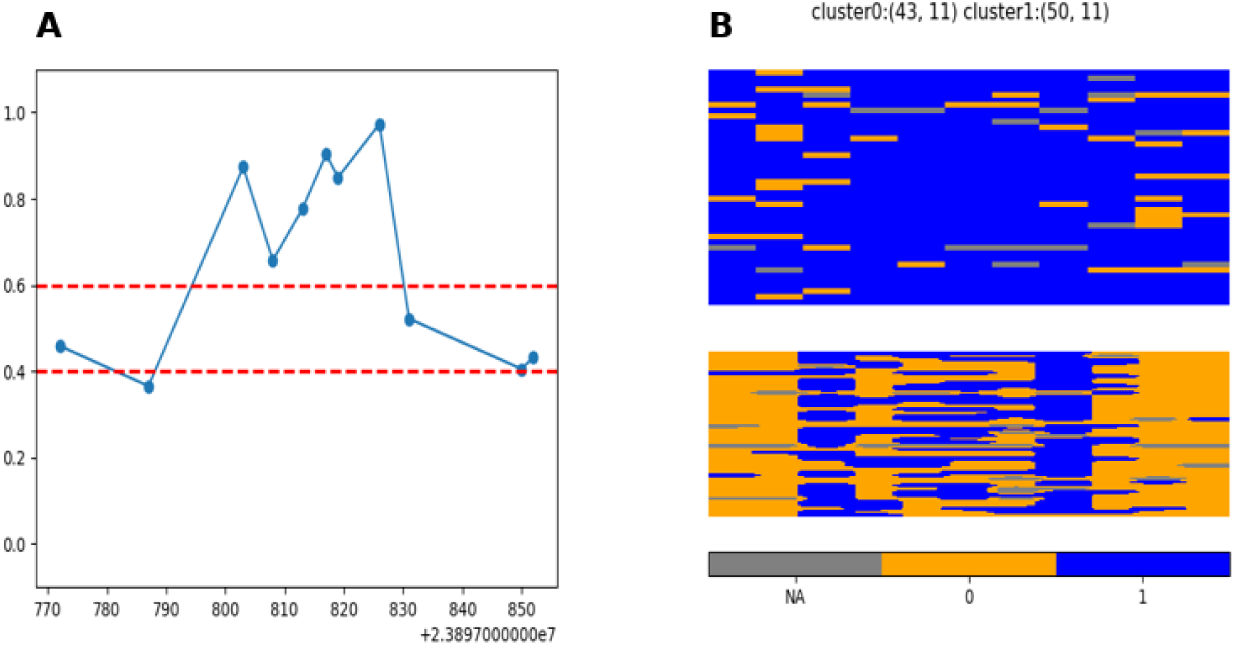
methylation in ICR region PWRN4(chr15:23897772-23897854) (A)methylation levels (B) clustered reads

**Figure S16:**
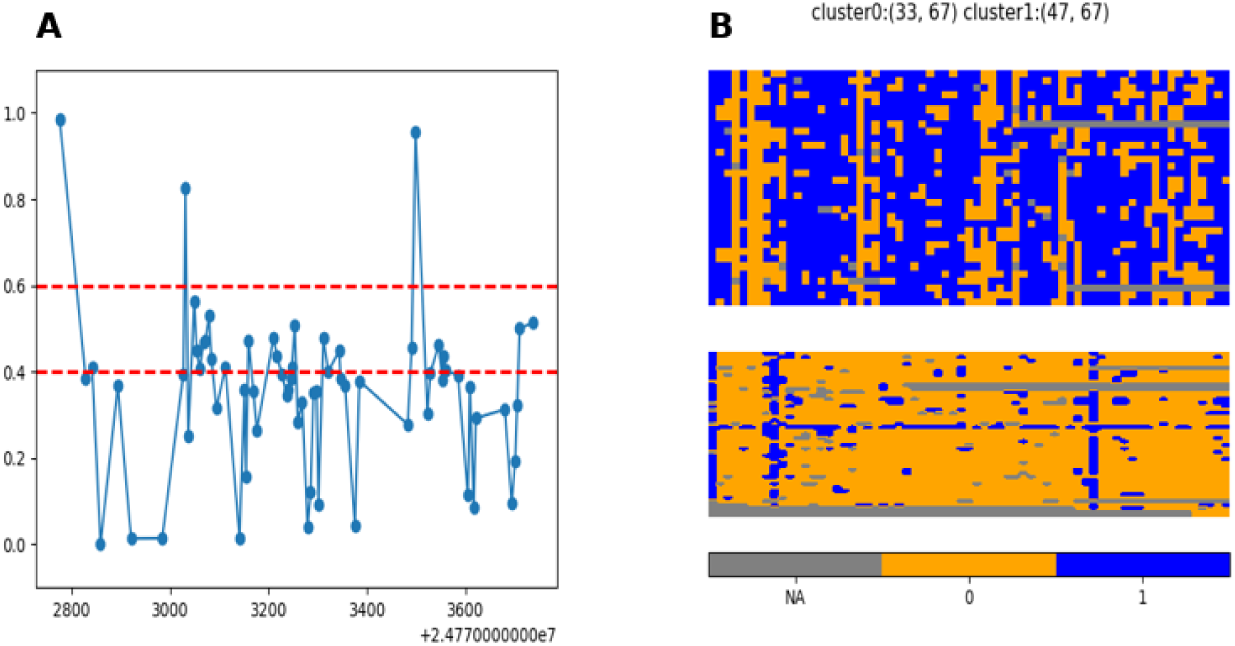
methylation in ICR region SNRPN_intragenicCpG40_(chr15:24772777-24773739) (A)methylation levels (B) clustered reads

**Figure S17:**
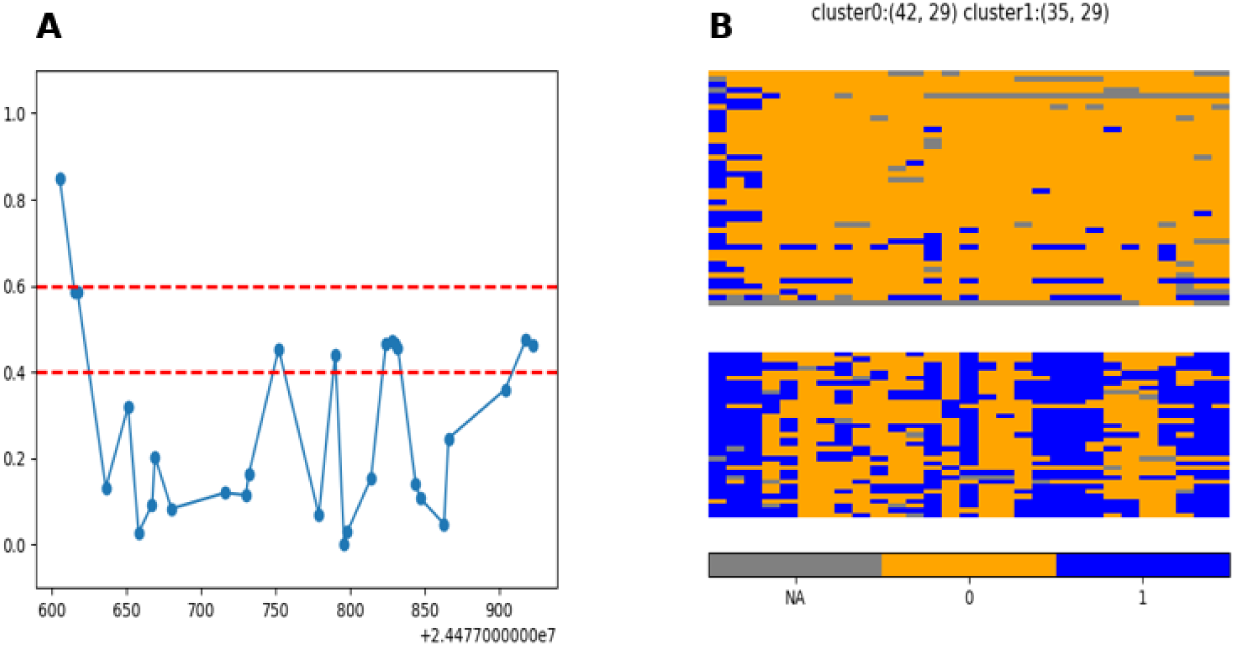
methylation in ICR region SNRPN_intragenicCpG30_(chr15:24477606-24477924) (A)methylation levels (B) clustered reads

**Figure S18:**
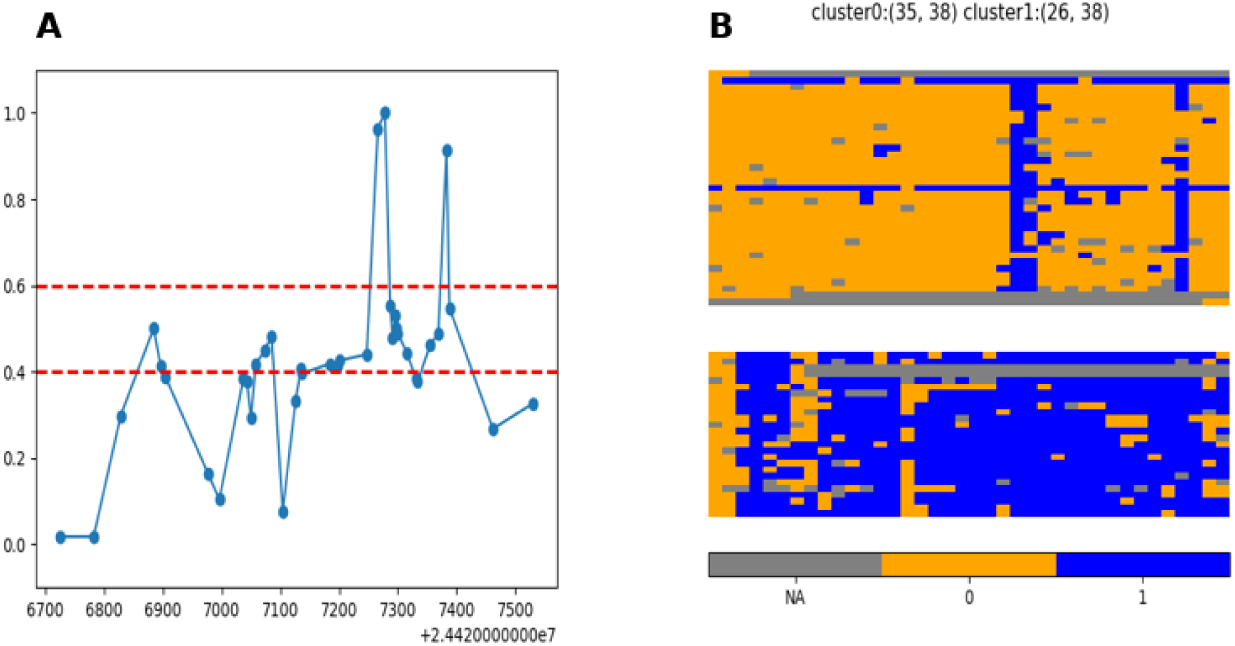
methylation in ICR region SNRPN_intragenicCpG29_(chr15:24426725-24427532) (A)methylation levels (B) clustered reads

**Figure S19:**
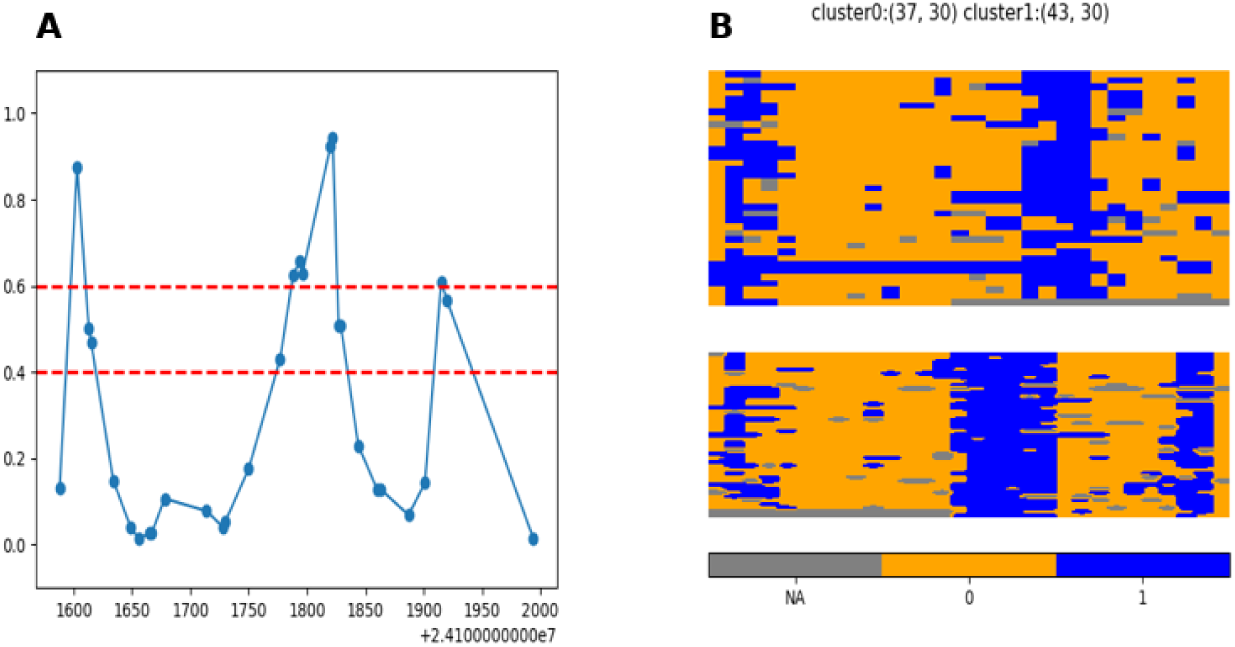
methylation in ICR region SNRPN_intragenicCpG32_(chr15:24101589-24101995) (A)methylation levels (B) clustered reads

**Figure S20:**
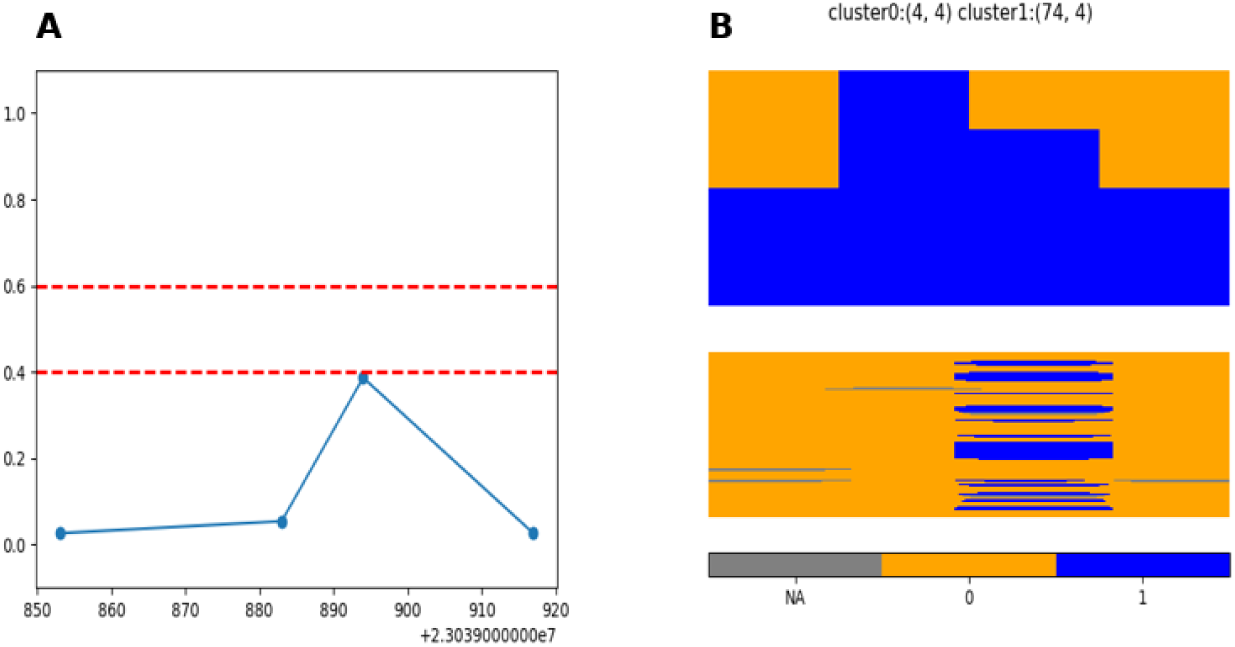
methylation in ICR region TUBGCP5(chr15:23039853-23039919) (A)methylation levels (B) clustered reads

**Figure S21:**
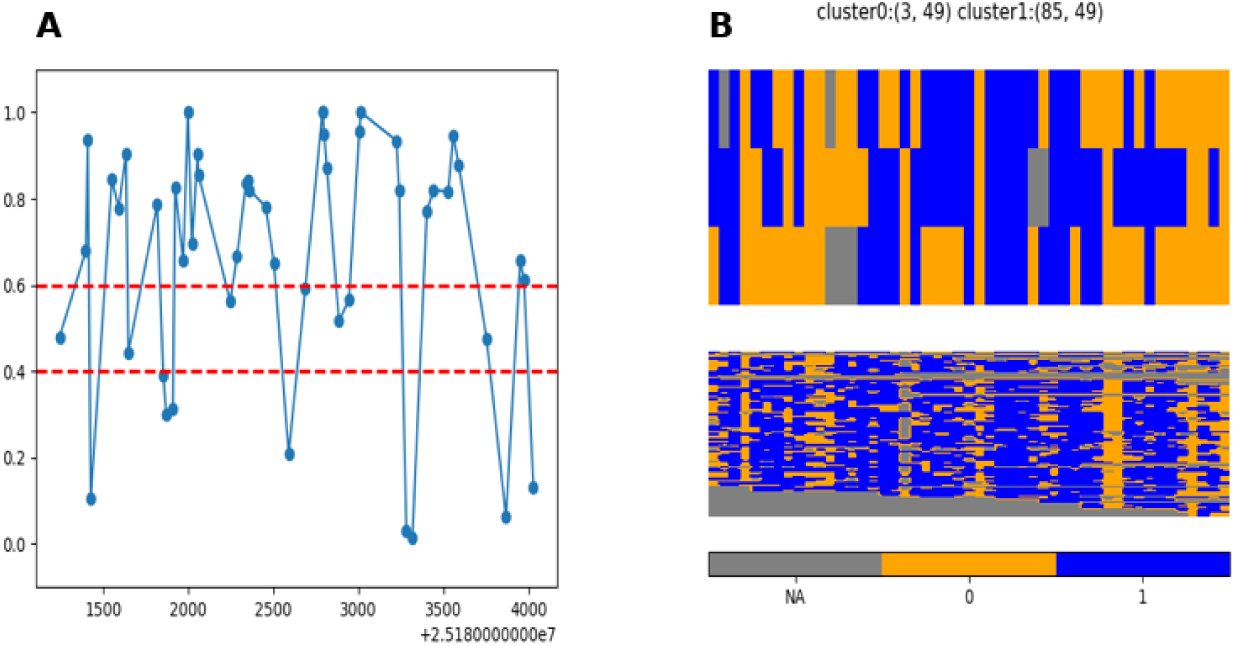
methylation in ICR region SNORD115Cluster(chr15:25181244-25184030) (A)methylation levels (B) clustered reads

**Figure S22:**
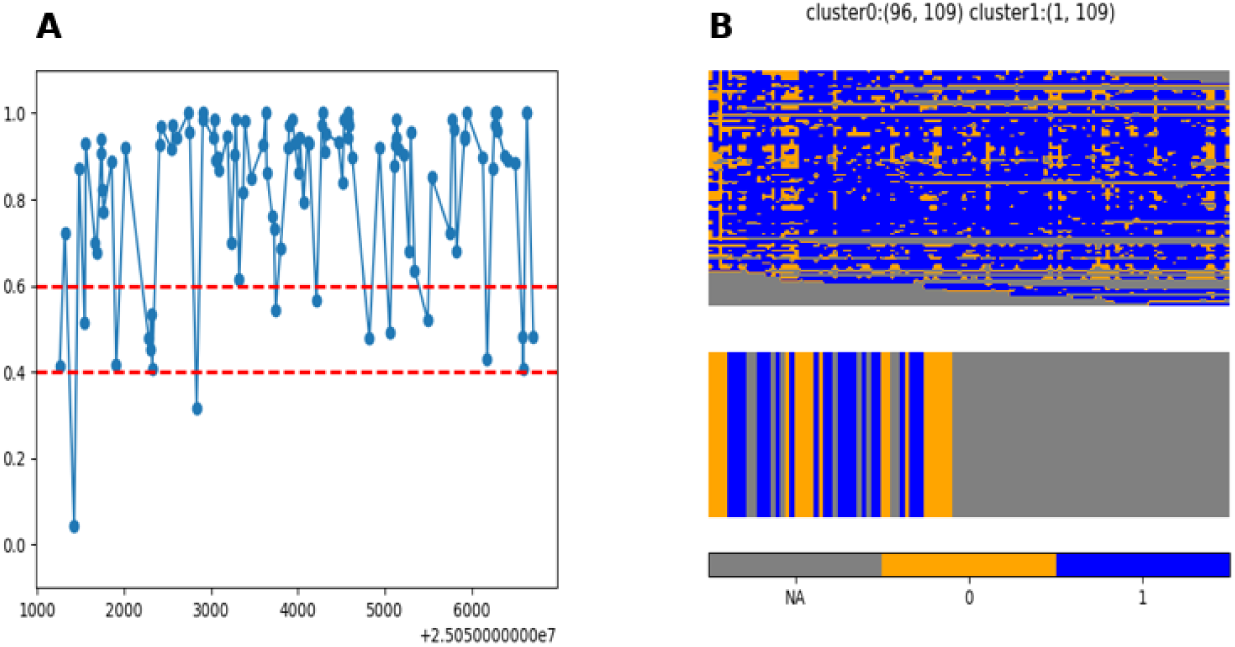
methylation in ICR region SNORD116Cluster(chr15:25051256-25056714) (A)methylation levels (B) clustered reads

**Figure S23:**
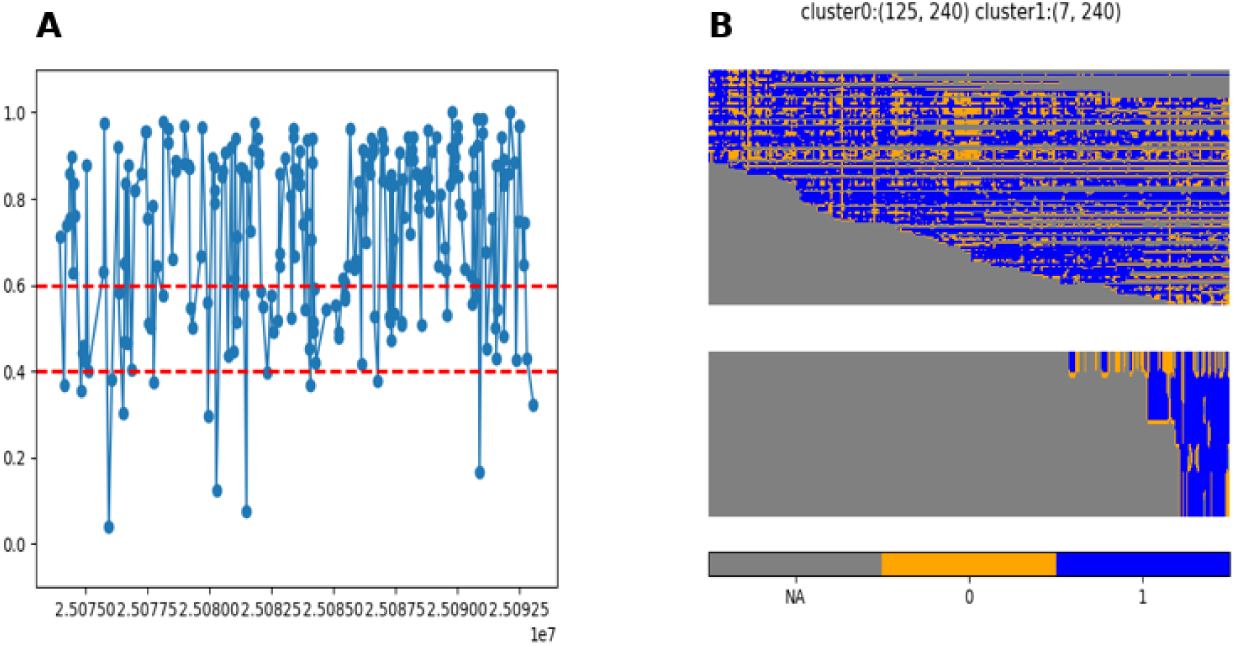
methylation in ICR region SNORD116Cluster(chr15:25073965-25093073) (A)methylation levels (B) clustered reads

**Figure S24:**
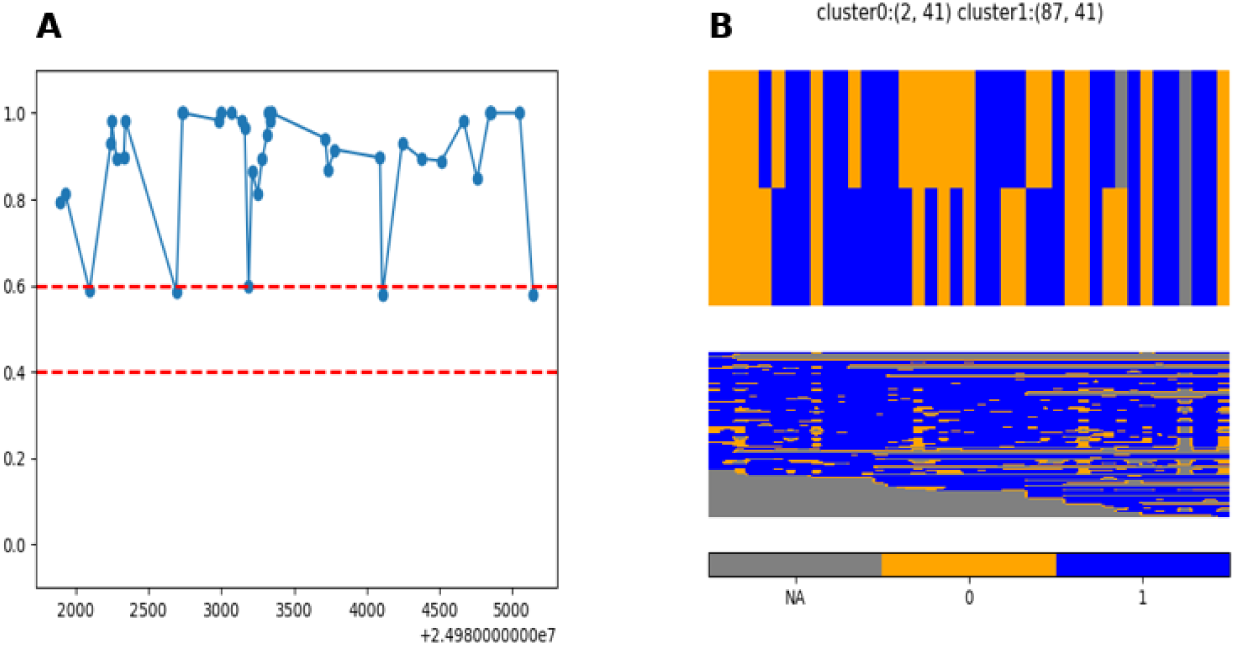
methylation in ICR region PWARSN,SNORD107,PWARSN,PWAR5(chr15:24981888-24985149) (A)methylation levels (B) clustered reads

### Fisher test on ICR regions

**Table 1:** Fisher test comparing the cluster assigned to each read by `asms` with the haplotag comouted ofor the same read by `whatshap`. low (significant) p-values indicate that haplotype assignment is correlated with cluster assignment.

|  |  |  |  |  |
| --- | --- | --- | --- | --- |
| chr15 | 98865267 | 98866421 | IGF1R | 2.271e-25 |
| chr15 | 24954857 | 24956829 | SNURF | 3.464e-24 |
| chr15 | 24856479 | 24856534 | SNRPN | 0.003582 |
| chr15 | 23647278 | 23648882 | MAGEL2 | 5.326e-16 |
| chr15 | 24847861 | 24848682 | SNRPN | 4.127e-13 |
| chr15 | 24877880 | 24878758 | SNRPN | 1.16e-16 |
| chr15 | 24823417 | 24824334 | SNRPN | 9.204e-07 |
| chr15 | 45022590 | 45022746 | SORD | 0.7008 |
| chr15 | 23686304 | 23687612 | NDN | 3.348e-16 |
| chr15 | 23561939 | 23567348 | MKRN3,MIR4508 | 7.508e-14 |
| chr15 | 23897772 | 23897854 | PWRN4 | 3.117e-21 |
| chr15 | 24772777 | 24773739 | SNRPN<sub>intragenicCpG40</sub> | 7.414e-09 |
| chr15 | 24477606 | 24477924 | SNRPN<sub>intragenicCpG30</sub> | 6.329e-15 |
| chr15 | 24426725 | 24427532 | SNRPN<sub>intragenicCpG29</sub> | 0.007143 |
| chr15 | 24101589 | 24101995 | SNRPN<sub>intragenicCpG32</sub> | 2.994e-15 |
| chr15 | 23039853 | 23039919 | TUBGCP5 | 0.03251 |
| chr15 | 25181244 | 25184030 | SNORD115Cluster | 0.1116 |
| chr15 | 25051256 | 25056714 | SNORD116Cluster | 1 |
| chr15 | 25073965 | 25093073 | SNORD116Cluster | 0.4397 |
| chr15 | 24981888 | 24985149 | PWARSN,SNORD107,PWARSN,PWAR5 | 0.2098 |

### scan-vcf

**Figure S25:**
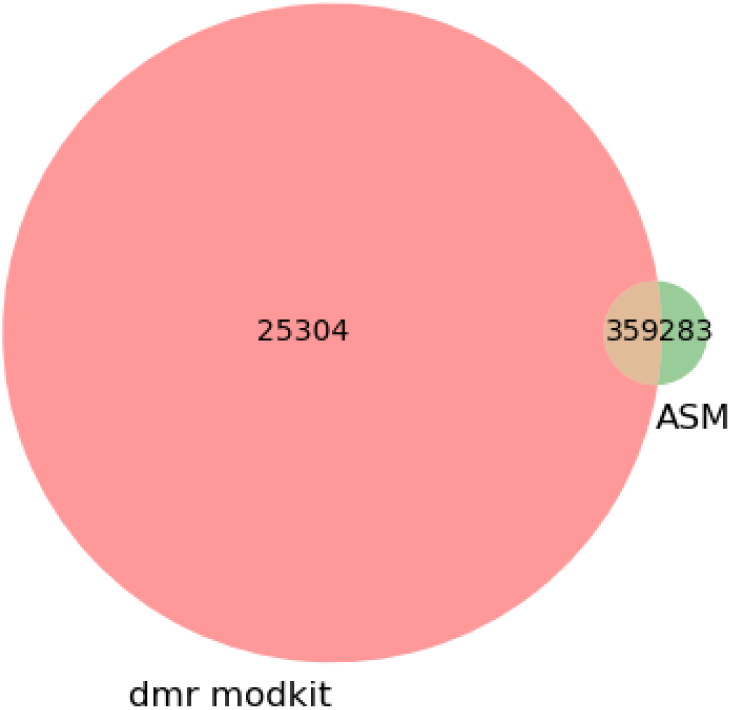
Venn diagram for the overlap between DMRs as found by modkit and ASM regions found by asms by looking in the neighbourhood of heterozygous loci.

